# Health professionals’ over estimation of knowledge on snakebite management, a threat to survival of snake bite victims - A cross-sectional study in Ghana

**DOI:** 10.1101/2020.09.02.280586

**Authors:** Evans Paul Kwame Ameade, Isaac Bonney, Evans Twumasi Boateng

## Abstract

**Background:** According to the World Health Organization, snakebites, a common occupational hazard in developing countries accounts an annual loss of between 81,000 and 139, 000 lives following 5 million of bites of which 2.7 million results in envenomation. Since snakebite associated morbidity and mortality is more prevalent in agriculture economies such as Ghana, health professionals should be optimally knowledgeable on how to manage incidence of snakebites. Lack of knowledge or overestimation of a professional’s knowledge can be affects heath delivery especially for emergency situations such as snakebites. The three (3) Tongu districts South Eastern Ghana which are rurally situated with agriculture as the major source of livelihood for their inhabitants are prone to snakebite incidence hence the need to assess whether the health professionals in these districts are well equipped by way of knowledge to handle such emergencies and whether they are able to rightly estimate their knowledge with regards to snakebite management.

**Methodology/Principal findings:** Data was collected using a *de novo* semi-structured questionnaire administered through google form whose link was sent via to 186 health workers made up of nurses, midwives, physician assistants, medical doctors, pharmacists, and pharmacy technicians. This data was analyzed using Statistical Package for the Social Sciences (SPSS) Version 25. Association between variables was determined using the appropriate tools where necessary, using a confidence interval of 95% and significance assumed when p ≤ 0.05. This study found male health workers significantly more knowledgeable about snakebite management (11.53±5.67 vrs 9.64±5.46; p = 0.022) but it was the females who overestimated their knowledge level (27.9% vrs 24.1%). The medical doctors exhibited the best knowledge on snakebite management with the registered general nurses least knowledgeable. Although most professionals overestimated their knowledge, the registered general nurses were the worst at that (53.7%). Overall knowledge of health care professionals on snakebite management was below average [10.60±5.62/22 (48.2%)] but previous in-service training and involvement in management of snakebite were associated with better knowledge. Respondents who had no previous training overestimated their knowledge level compared to those who had some post qualification training on snakebite management (7.5% vrs 38.1%). Greatest knowledge deficit of respondents was on the management of ASV associated adverse reactions.

**Conclusion:** Health workers in rural Ghana overestimated their knowledge about snakebite management although their knowledge was low. Training schools therefore need to incorporate snakebite management in their curriculum and health authorities should also expose health workers to more in-service training on this neglected tropical disease.

**Author summary:** World Health Organization estimates that every year between 81,000 and 139,000 die due snake bites across the world. Mismanagement of snakebites can result in increased disabilities and death if not handled by knowledgeable health workers. This study assessed if various categories of health workers made up of professionals from the medical, pharmaceutical and nursing categories in the three neighbouring Tongu districts in Ghana have the appropriate level of knowledge on snakebite management. Using a newly developed questionnaire, data was collected from the respondents using google forms sent to their WhatsApp platforms. Data was then analyzed using Statistical Package for the Social Sciences (SPSS) Version 25. Results were presented in the form of tables and association between the variables also determined. The level of knowledge of sampled health workers on snakebite was below average especially among the nursing professionals. However, those who had some previous post qualification training on snakebite management exhibited a significant superior knowledge and least overestimated their knowledge hence policy makers should through workshops equip health workers especially the nurses on snakebites so that rural dwellers whose health care needs are mainly attended to by nurses can be better managed when they suffer snakebites.

## INTRODUCTION

Snakes which belongs to the class of animals called the reptiles can be found in all places except in Antarctica, Iceland, Ireland, Greenland, New Zealand, Cape Verde in West Africa, Siberia area in Russia, some parts of Argentina, Chile, Finland as well as some small nations in the Pacific Ocean such as Tuvalu and Nauru [1]. It is estimated that there are more than 3,700 species of snakes on earth [2]. As snakes also makes efforts to survive in the ecosystem, there are bound to come into conflict with humans and mostly as a defensive mechanism some of them bite. This human-snake conflict is estimated to results in between 4.5 and 5.4 million snakebites annually [3]. It is estimated that about 600 snakes whenever they bite, they inject toxins substances referred to as venoms into their victims hence they are classified as being venomous while the vast majority are non-venomous [4]. The number of persons bitten by venomous snakes cannot be exactly known but it is believed that 1.8 to 2.7 million people globally suffer the effects of their bites out of which 81,000 to 138,000 of victims die although the mortality would have been higher had it not been because about 50% of venomous snakebites do not lead to envenoming [4, 5]. Notwithstanding this high level of snakebite incidence, reports across the world found that quite a number of victims seek remedies from traditional medicine practitioners than hospitals. A study in India found that only 22.2% of snakebite victims report at the hospitals [6]. Two hospital-based surveys in Nigeria and Ghana reported snakebite incidence of 465 per 100,000 and 92 per 100,00 respectively [7, 8]. Mortality and morbidity associated with snakebites for those who report at the hospital can be determined by the level of management by the health care professionals which will depend on how knowledgeable or skillful they are on snakebite management. There is paucity of study on assessment of the knowledge of healthcare professionals on the management of snakebites in Ghana hence the need to undertake this study in three rural districts of Ghana in the coastal savanna eco-zone.

## METHOD

### Study setting

The study areas are selected health facilities South, Central and North Tongu districts of the Volta region of Ghana. The facilities in the South Tongu district were the District Hospital and Comboni Catholic Hospital both located at Sogakofe; Health Centres at Tefle Kpotame and Adutor and the Agbakofe and Sasekofe Community-based Health Planning and Service (CHPS) zones. CHPS zones are the lowest level of health care system in Ghana for the provision of primary health care to those in rural Ghana. For the North Tongu District, Battor Catholic Hospital and Volo Health Centre were the sites for the study while the Central Tongu District had Mafi Adidome Hospital as well as Mafi Kumase and Mafi Dove Health Centres as the study sites. The total population of these three Tongu districts in Ghana’s 2010 National population census was 237,138 [9]. Inhabitants of these districts (Figure 1) whose main occupation are agriculture related, speak mainly the Tongu dialect of the Ewe language.

**Figure 1:**
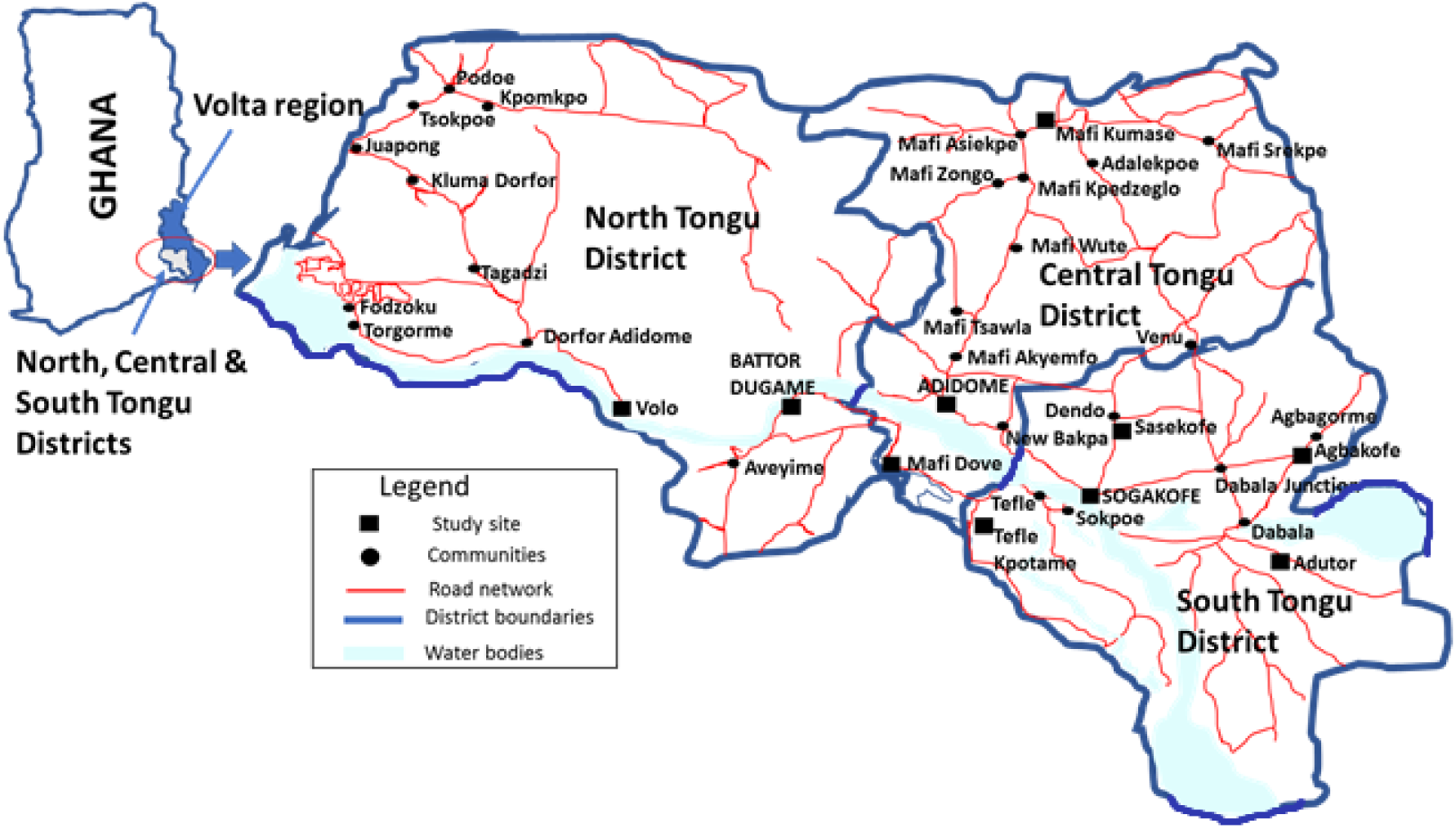
The map of Ghana and the study location, the South, Central and Tongu Districts of the Volta region of Ghana.

### Study design

A cross-sectional study design was applied in this study which was conducted within the months of May and June, 2019.

### Study population

The study population were health care providers namely; pharmacists, physician/medical assistants, medical doctors, pharmacy technicians as well as midwives and nurses of various categories who work in hospitals, health centres and CHPS compounds in the study area.

### Study sample size determination

The sample size for this study was calculated using the Cochran formula, 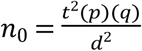

Where t (selected alpha level of .025 in each tail) = 1.96; d, (acceptable margin of error) = 0.05; With an estimated overall knowledge of health care providers in the study area on management of snakebites as 50%, p (the estimated proportion of an attribute that is present in the population) = 0.5 hence q is 1-p = 0.5.

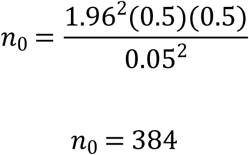

Since *n*_0_ of 384 exceeds the 5% of the eligible study population of 537 excluding the 20 involved in the pre-testing of the questionnaire (537 × 0.05 = 26.9), Cochran correction formula can be used to obtain adjusted sample size *n*_1_

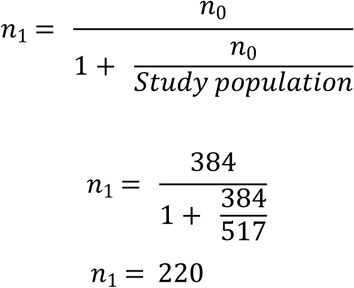

With as expected response rate of 90%, the final actual sample size for the study was 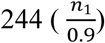.

At the end of the study period, responses from 186 individuals were successfully received resulting in a response rate of 76.2% (186/244*100).

### Sampling procedure

Efforts were made to take a census sample of all the pharmacists (5), physician/medical assistants (26), medical doctors (17), and pharmacy technicians (6) because of their small numbers in the selected health facilities. However, for the nurses and midwives who were about 483, convenience sampling technique was applied to select the respondents. For the category of health professionals that census technique was applied, they were met in person or spoken to on their mobile phone and the link of the questionnaire was sent to the WhatsApp pages of those willing to partake in the study. All nurses and midwives working in Health centres and CHPS zones which are the lowest level of health care in Ghana, were invited to partake in the study after a visitation by a member of the research team. For the respondents fron the hospitals, invitation was extended to those who were at the facility at the time of the visit of the research team. Some off-duty nurses and midwives were gotten in to participate in the study by their colleagues who the researchers had earlier met and enrolled into the study.

#### Data Collection Instrument and Technique

A de novo self-administered semi-structured questionnaire was designed and then converted into google form. The questions were formulated based on information obtained from the 2016 edition of the WHO Guidelines for the management of snakebites, WHO Regional Office for South-East Asia. The questionnaire was piloted among twenty(20) health workers from the study area who were subsequently excluded during the actual data collection. The research team performed a face validity of the questionnaire and also deleted or reframed questions that were ambiguous so as to ensure their clarity. Reliability test of the questionnaire was also performed using Microsoft Excel which gave a Cronbach alpha of 0.7 which made the questionnaire acceptable for the study. The questionnaire consisted of thirty-one questions of which six were on respondents’ sociodemographic characteristics, and another nine related to previous training and management as well of their level of confidence in the management of snakebites. The remainder fifteen questions assessed respondents’ knowledge about snakebite management. The questionnaire was administered through the WhatsApp accounts of the respondents using the link https://forms.gle/iV5NtKzdjbg5LTSc9. Follow up text messages were sent and calls made to the respondents to remind them of the need to complete and submit the questionnaire.

### Data measurement

Comparing the answers provided by the respondents with those from literature sources, the knowledge of the health professionals on snakebite management was assessed. For questions that the respondents had the option of choosing the most appropriate answer, a score of 1 mark was allocated. Choosing a wrong or an ‘I don’t know’ option attracts zero mark. The total score for open-ended questions depends on the maximum number of answers required to be provided hence a question that required the provision of four answers scores 4 marks if all the respondent’s answers are considered appropriate. The total maximum score which assessed the respondent’s knowledge on snakebite management was 22. In comparing the perceived and actual knowledge scores of respondents on snakebite management, the total score of actual knowledge of each respondent was converted to 10 because respondents stated their perceived knowledge on snakebite management with values ranging between 0 and 10 with 0 indicating absolute lack of knowledge while 10, for the most excellent level of knowledge.

### Statistical analysis

Descriptive data was presented in tables indicating frequencies and percentages of the variables and questions used for the assessment. Association between variables was also determined using One-Way ANOVA at a confidence interval of 95%. Assessment of the level of over or under estimation or exaggeration of respondent’s was measured by comparing respondents perceived knowledge and their actual knowledge score using paired sample test and pair sample correlation. Significance was assumed when p ≤ 0.05.

#### 3.11 Ethical Consideration

The ethics committee of the School of Medicine and Health Sciences of the University for Development Studies, Tamale provided ethical clearance for this study. Again, the preamble on the questionnaire explained the purpose of the research and stating clearly that submitting the form after completion is indicative of giving consent. To ensure confidentiality, the names of the respondents were not required. Clearances were also obtained from the District Health Directorates as well as the administrative heads of the various health facilities from which the data were collected.

## RESULTS

### Sociodemographic characteristics of respondents

The number of male and females who took part in the study was almost equal (51.1% vrs 48.9%) but those between ages 30 and 39 years were in the majority, 98 (52.7%). Again, majority of respondents were health workers in hospitals, 146 (78.5%) and had worked for less than 5 years, 112 (60.2%). Most respondents were from the South Tongu District, 87 (46.8%) and were registered general nurses, 80 (43.0%) but the health profession least represented were the pharmacists, 4 (2.2%). Table 1 shows the sociodemographic characteristics of respondents in this study.

**Table 1:** Socio-demographic characteristics of respondents.

| Variable | Subgroup | Frequency | Percentage |
| --- | --- | --- | --- |
| Sex | Male | 95 | 51.1 |
|  | Female | 91 | 48.9 |
| Age (years) | 20-29 | 81 | 43.5 |
|  | 30-39 | 98 | 52.7 |
|  | >39 | 7 | 3.7 |
| District | South Tongu | 87 | 46.8 |
|  | Central Tongu | 45 | 24.2 |
|  | North Tongu | 54 | 29 |
| Number of years of practice (years) | < 5 | 112 | 60.2 |
|  | 5 – 10 | 71 | 38.2 |
|  | >10 | 3 | 1.6 |
| Level of health facility | CHPS zones | 15 | 8.1 |
|  | Health Centre | 25 | 13.4 |
|  | Hospital | 146 | 78.5 |
| Profession category | Registered General Nurse | 80 | 43.0 |
|  | Enrolled/Community Nurse | 37 | 26.3 |
|  | Midwife | 15 | 8.1 |
|  | Medical officer | 14 | 7.5 |
|  | Pharmacy Technician | 5 | 2.7 |
|  | Pharmacist | 4 | 2.2 |
|  | Physician/Medical assistant | 19 | 10.2 |

### Training on and management of snakebite

Table 2 presents the record of post qualification training on snakebite management and management history of respondents. Although, those who had ever been provided training on snakebite management since they started practicing as healthcare professionals were in the minority, 57 (30.6%), majority of the respondents, 154 (82.8%) had ever been involved in the management of snakebite victims in their facilities. For those who had no post qualification formal training on snakebite management, most, 53 (40.2%) had snakebite management skills from their senior colleagues with a lesser number, 32 (24.2%) acquiring their knowledge by reading materials from the internet and books. For the first half of the year 2019, most, 92 (49.5%) respondents who had ever managed snakebite cases had taken care of between 1 and 5 victims. Although majority, 160 (86.0) will triage snakebite as emergency, most respondents, 91 (48.9%) do not think their health facilities have all the resources for optimal management of snakebites. The major limitation against the management of snakebite for those who think their health facilities cannot manage snakebites adequately is the unavailability of anti-snake venom,77 (86.5%) although majority of respondents, 139 (73.7%) of all respondent said their health facilities have protocols for the management of snakebites. Most respondents, 79 (43.6%) were fairly confident about their ability to manage snakebite victims.

**Table 2:** Training on and management of snakebite incidence by respondents.

| Variable | Subgroup | Frequency | Percentage |
| --- | --- | --- | --- |
| Ever managed snakebite | Yes | 154 | 82.8 |
|  | No | 32 | 17.2 |
| Ever been trained on snakebite management after school? | Yes | 57 | 30.6 |
|  | No | 129 | 69.4 |
| If not trained, how was skill acquired? (n = 132) | Learning from senior colleagues on the job | 53 | 40.2 |
|  | Knowledge and skills obtained in school | 47 | 35.6 |
|  | Self-education on the internet or in text books | 32 | 24.2 |
| Number of snakebites managed half year (January to June, 2019) | 0 | 77 | 41.4 |
|  | 1-5 | 92 | 49.5 |
|  | 6-10 | 12 | 6.5 |
|  | >10 | 5 | 2.7 |
| How to you triage snakebite? | Emergency | 160 | 86.0 |
|  | Urgent | 24 | 12.9 |
|  | Don't know | 2 | 1.1 |
| Does your health facility have what it takes to manage snakebites? | Yes | 90 | 48.4 |
|  | No | 91 | 48.9 |
|  | I don't know | 5 | 2.7 |
| Reasons for which your health facility unable to manage snakebites (n = 89) | Lack or inadequate Anti Snake venom | 77 | 86.5 |
|  | Lack of other logistics | 10 | 11.2 |
|  | Inadequate qualified staff | 2 | 2.2 |
| Does your hospital have snakebite management protocol? | Yes | 137 | 73.7 |
|  | No | 34 | 18.3 |
|  | I don't know | 15 | 8.1 |
| How confident are you about snakebite management (n = 181) | Not confident | 11 | 6.1 |
|  | Fairly confident | 79 | 43.6 |
|  | Confident | 78 | 43.1 |
|  | Very confident | 13 | 7.2 |

### Knowledge of respondents on snakebite management

Table 3 shows the level of knowledge of respondents on snakebite management. The top five best answered questions on snakebite management were; Antivenoms being the only specific antidotes in the management of snake bites by venomous snakes [0.92±0.273 (92.0%)], injecting ASV intramuscularly is as not as effective as using the intravenous route [0.81±0.39 (81.2%)], the 20 minutes whole blood count test (20MWBCT) being the first recommended test for a suspected snakebite victim to determine envenoming [0.73±0.447 (73.0%)], stating correctly three adverse reactions a patient given anti-snake venom (ASV) may experience [2.01±1.24 (67.0%)] and antivenoms need not be given to all persons suspected of snakebite [0.67±0.47 (67.0%)]. The bottom five areas of least knowledge about snakebite management by the respondents were; ASV being useful for months and years after the labelled expiry date [0.12±0.32 (12.0%)], intramuscular route being the most appropriate for administering first choice drug used for managing adverse reaction caused by ASV [0.19±0.40 (19.4%)], Adrenaline being the first choice in the management of adverse reactions caused by ASV rather than hydrocortisone which majority, 96 (51.6%) wrongly indicated [0.22±0.42 (22.0%)], a snake bite patient reporting to a facility with a tourniquet applied to the affected limb must be told it is not appropriate, but informed that the tourniquet will not be removed until anti-snake venom is injected [0.22±0.42 (22.0%)] and correctly stating any important biochemical test required in snakebite management [0.31±0.464 (31.2%)]. The overall knowledge score of the respondents on snakebite management was 10.60±5.62 over 22 which is equivalent to 48.2%.

**Table 3:**
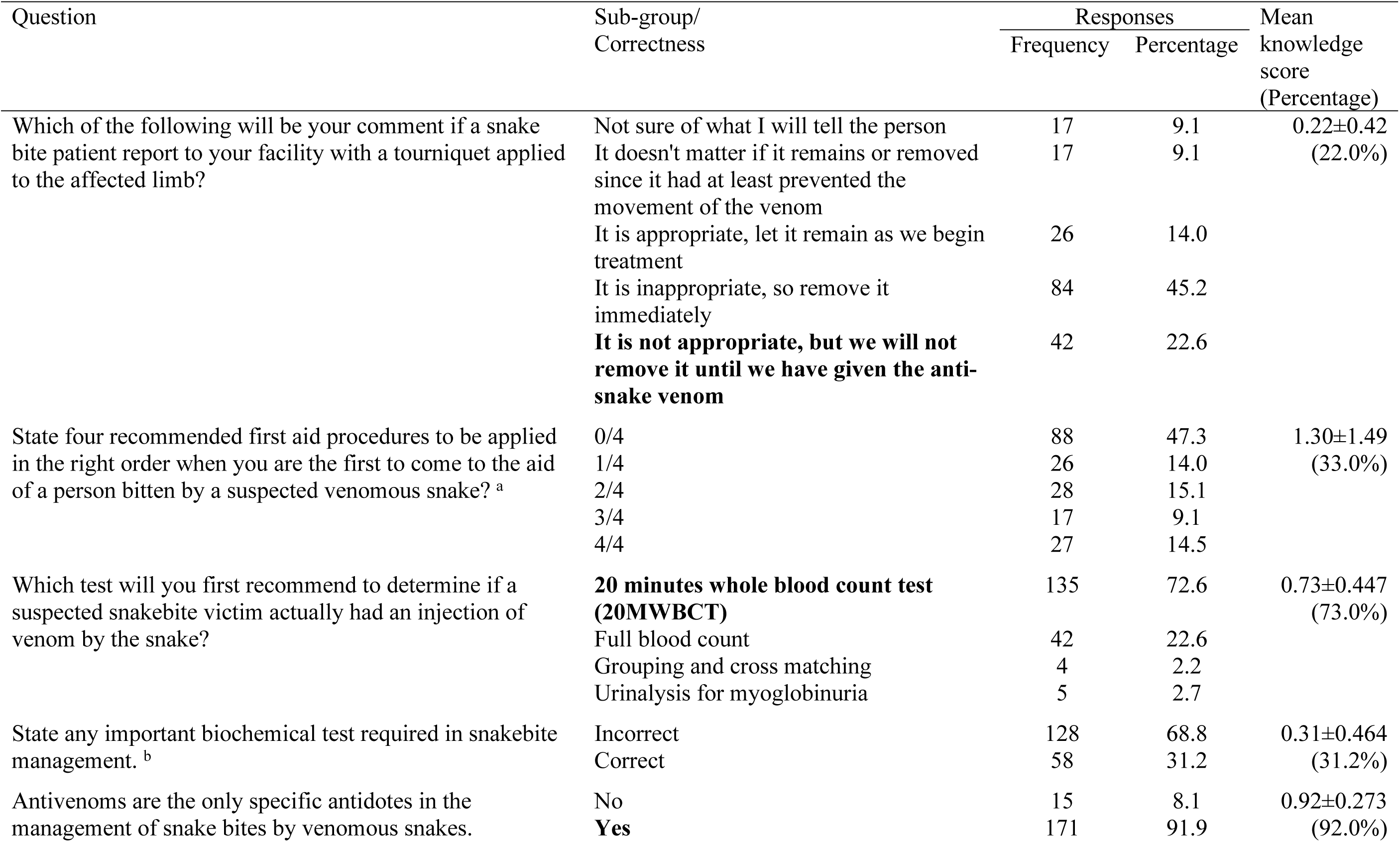
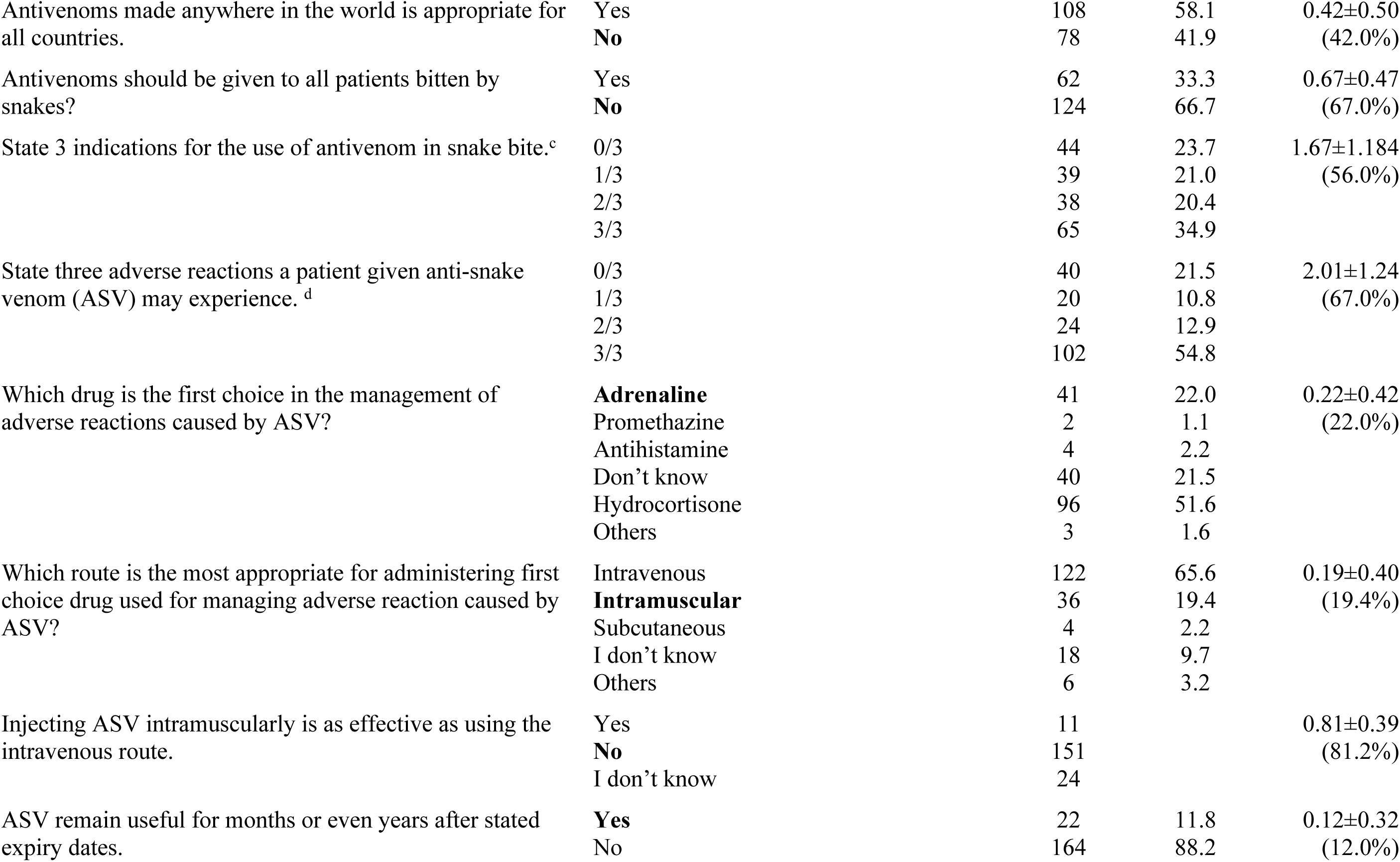
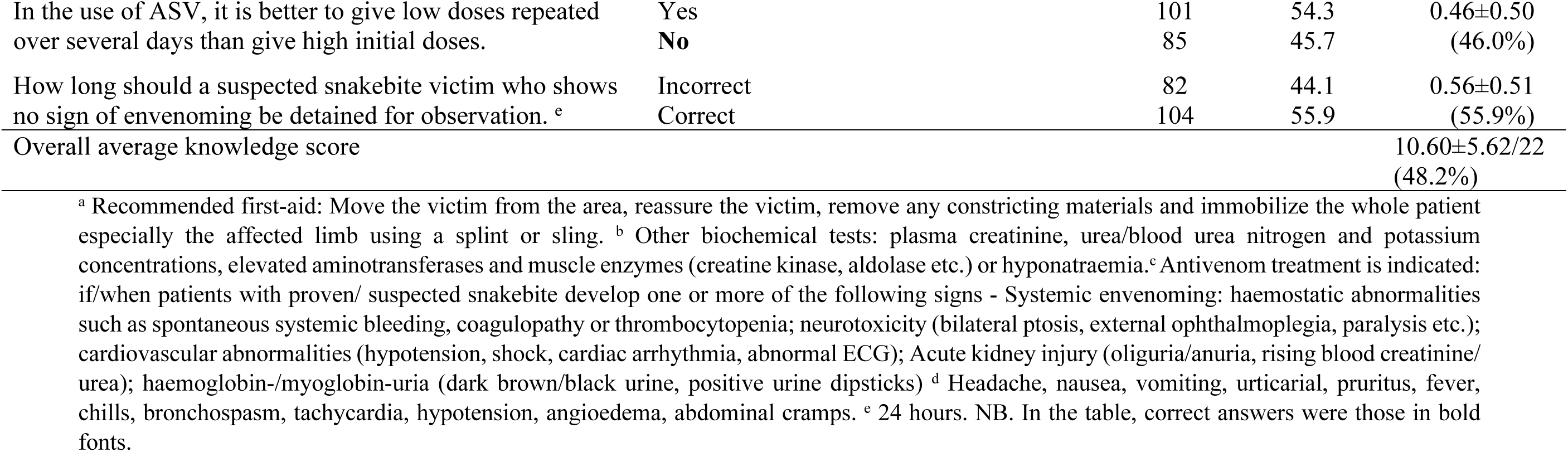
Knowledge of respondents on snakebite management.

### Association between socio-demographic characteristics and knowledge on snakebite management

Table 4 shows the association between socio-demographic characteristics and knowledge on snakebite management. Male respondents were significantly more knowledgeable about snakebite management than females (11.53±5.67 vrs 9.64±5.46; p = 0.022) so also were those who had some previous training on snakebite management than those who were not provided any other form of in-service training (14.14±5.90 vrs 9.04±4.75; p <0.001). Previous experience on snakebite management provides significantly better knowledge on snakebite management than one who had never been involved in the management of snakebite (5.17±2.47 vrs 3.15±2.38; p <0.001). Respondents working at CHPS zones scored best (14.47±5.48) followed by those at health centres (12.72±6.88) with those at hospitals being the least knowledgeable (9.84±5.18) on snakebite management with the differences in knowledge being statistically significant (p <0.001). There were significant differences in knowledge among respondents based on their district of practice (p = 0.003) with those from the North Tongu District scoring the highest (11.94±5.95), closely followed by those in the Central Tongu District (11.84±5.93) whereas the South Tongu district respondents scored the least (9.84±5.18). There was a significant difference on knowledge on snakebite among the various categories of healthcare professionals (p = 0.031) with the medical doctors obtaining the best mean score of 13.71±6.50 followed by the Pharmacy technician (13.60±6.07) and the pharmacist (13.50±7.77) but the registered general nurses were the worst performers (9.11±4.63). Further grouping of the various categories of health workers based on their core duties found the prescribers being the most significantly knowledgeable group (13.56±6.41; p = 0.017) with the nursing and midwifery group scoring the least (9.98±5.31).

**Table 4:**
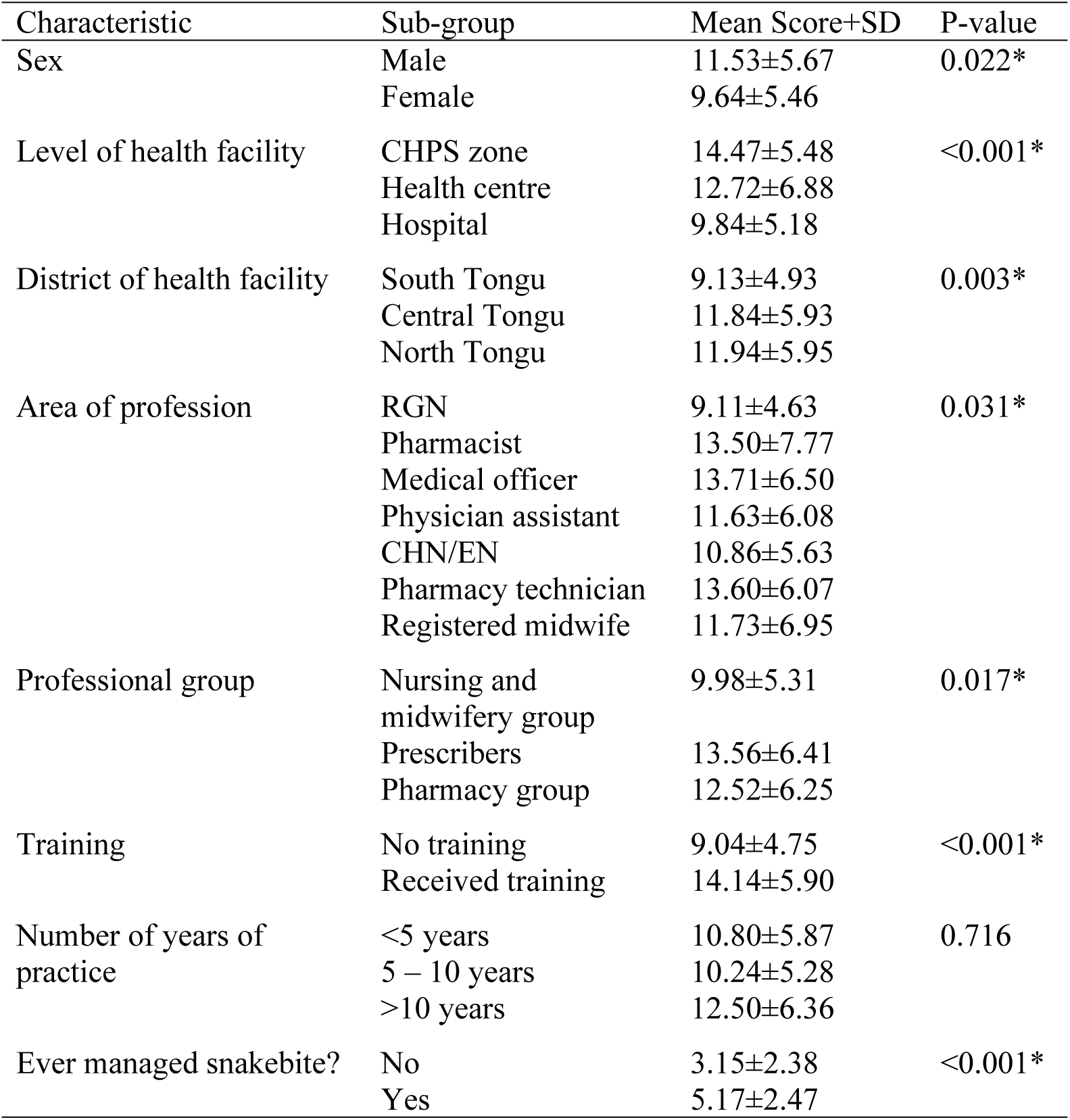
Association between socio-demographic characteristics and actual knowledge scores on snakebite management.

### Comparison between the perceived and actual knowledge scores of respondents on snakebite management and

Table 5 shows the comparison between the perceived and actual knowledge scores of respondents on snakebite management against their sociodemographic characteristics. There was significant difference between the mean actual and perceived knowledge scores on snakebite management for both male (p < 0.001) and female (p < 0.001) respondents but the females had greater exaggerated confidence than the males (24.9% vrs 24.1%). For both the male (p = 0.008) and the female (p = 0.003) respondents, there was a small but significant positive correlation (≈ 0.3) between their perceived and actual knowledge scores. All the age categories had exaggerated snakebite management knowledge scores > 20.0% but those above 39 years had the highest score difference of 43.5%. However, it was only age groups 20 to 29 and 30 to 39 that had the differences between their perceived and actual knowledge scores being statistically significant (p < 0.001). Again, although all age categories had weak positive correlation between the mean perceived and actual scores, it was only those between 30 and 39 that that had a significant correlation (p < 0.001). Whereas health workers in the lowest level of Ghana’s healthcare system, the CHPS zones significantly underestimated their knowledge on snakebite management (−25.4%; p = 0.005), their colleagues in the hospitals significantly over estimated their knowledge (37.4%; = < 0.001). It was only respondents in the hospitals that had a moderate but significant correlation between their actual and perceived knowledge on snakebite management. Health workers in the South Tongu District had the highest significant over exaggeration of knowledge on snakebite management (46.6%; p < 0.001) as well as a significant and strong correlation between perceived and actual knowledge scores (r = 0.5; p < 0.001). All the categories of the number of years of practice had an over exaggerated score but those practicing between 5 and 10 years recorded the highest significant difference score (38.4%; p < 0.001) while those who had worked for less than a year had the lowest significant difference score (17.8%; p = 0.001). Again, whereas those who had practiced for less than 5 years had a moderate but significant correlation (r = 0.3; p = 0.01) between the perceived and actual mean knowledge score, those who had worked between 5 and 10 years had a strong and significant correlation (r = 0.5; p < 0.001). There was over exaggeration of knowledge on snakebite management whether respondents had over had any form of on-the-job training or not (7.5% vrs 38.1%) but those who had no training had a significant exaggeration (p < 0.001) and also a moderate but significant correlation between their actual and perceived mean snakebite management scores (r = 0.4; p < 0.001). Only the pharmacists (−2.3%), midwives (−12.9%) and the pharmacy technician (−22.2%) under estimated their knowledge on snakebite management. Other healthcare workers such as medical doctors (21.5%), physician/medical assistants (24.4%) and registered general nurses (53.7%) over exaggerated their knowledge by more than 20.0% but it was only the registered general nurses (p < 0.001) and the physician/medical assistants (p = 0.032) that the differences between their perceived and actual knowledge scores were significantly different. Whereas, there was a moderate but significant correlation between perceived and actual knowledge scores of the registered general nurses (r = 0.4; p < 0.001), that of the physician/medical assistants was a strongly significant correlation (r = 0.5; p = 0.037). Regrouping of the health workers categories based on their core functions found significant differences between perceived and actual knowledge score for the nursing/midwifery group (29.6%; p < 0.001) and the prescribers (23%; p = 0.005) as well as a significant and moderate correlation (r = 0.3; p = 0.002) for the nursing/midwifery group but a significant and strong correction for the prescribers (r = 0.5; 0.005). Both respondents who had in-service training and those without over exaggerated their knowledge (26.4% vrs 18.9%) but it was only those who had been trained who shows a significant difference between their perceived and actual knowledge scores. Whereas the correlation between the scores were for those without training was strong and significant (r = 0.5; p = 0.011), those who had training was significant but weak (r = 0.2; p = 0.042). All the forms of verbal declaration of confidence in snakebite management showed exaggeration of knowledge with those who stated they had no confidence scoring the least difference (6.6%) which was not significant and those who stated they were very confident recording a significantly large difference between actual and perceived knowledge scores (52.7%; p < 0.001).

**Table 5:** Comparing means of respondents perceived and actual knowledge of snakebite management according to their sociodemographic characteristics.

| Variable | Subgroup | Frequency | Paired sample statistics |  | Paired samples test |  | Paired samples correlation |  |
| --- | --- | --- | --- | --- | --- | --- | --- | --- |
|  |  |  | Mean<br>NPS±SD | Mean<br>AKS±SD | Difference (%) | p value | r | P value |
| Sex | Male | 93 | 6.60±1.62 | 5.32±2.54 | 1.28±2.62 (24.1) | <0.001* | 0.3 | 0.008* |
|  | Female | 88 | 5.73±1.82 | 4.48±2.46 | 1.25±2.55 (27.9) | <0.001* | 0.3 | 0.003* |
| Age | 20-29 | 78 | 5.68±1.49 | 4.52±2.36 | 1.16±2.60 (25.7) | <0.001* | 0.1 | 0.200 |
|  | 30-39 | 96 | 6.49±1.90 | 5.22±2.63 | 1.27±2.57 (24.3) | <0.001* | 0.4 | <0.001* |
|  | >39 | 7 | 7.36±1.18 | 5.13±2.63 | 2.23±2.57 (43.5) | 0.062 | 0.3 | 0.550 |
| Level of facility | CHPS | 14 | 5.21±0.70 | 6.98±2.01 | -1.77±1.94 (-25.4) | 0.005* | 0.3 | 0.346 |
|  | Health Centre | 21 | 5.86±1.35 | 6.28±3.06 | -0.42±3.04 (-6.7) | 0.534 | 0.2 | 0.298 |
|  | Hospital | 127 | 6.24±1.88 | 4.54±2.40 | 1.70±2.27 (37.4) | <0.001* | 0.5 | <0.001* |
| District | South Tongu | 84 | 6.23±2.03 | 4.25±2.21 | 1.98±2.12 (46.6) | <0.001* | 0.5 | <0.001* |
|  | Central Tongu | 44 | 6.21±1.53 | 5.47±2.67 | 0.74±2.80 (13.5) | 0.087 | 0.2 | 0.197 |
|  | North Tongu | 53 | 6.06±1.51 | 5.51±2.67 | 0.55±2.78 (10.0) | 0.156 | 0.2 | 0.137 |
| Number of years of practice (years) | <5 | 109 | 5.90±1.54 | 5.01±2.64 | 0.89±2.71 (17.8) | 0.001* | 0.3 | 0.010* |
|  | 5-10 | 69 | 6.56±2.03 | 4.74±2.38 | 1.82±2.30 (38.4) | <0.001* | 0.5 | <0.001* |
|  | >10 | 3 | 7.33±1.53 | 5.91±2.08 | 1.42±1.87 (24.0) | 0.318 | 0.5 | 0.667 |
| Ever had training on Snakebite management? | No | 125 | 5.80±1.77 | 4.20±2.14 | 1.60±2.24 (38.1) | <0.001* | 0.4 | <0.001* |
|  | Yes | 56 | 7.01±1.45 | 6.52±2.62 | 0.49±3.09 (7.5) | 0.239 | -0.1 | 0.558 |
| Profession | RGN | 78 | 6.47±1.93 | 4.21±2.08 | 2.26±2.16 (53.7) | <0.001* | 0.4 | <0.001* |
|  | Pharmacist | 4 | 6.00±2.94 | 6.14±3.53 | -0.14±2.18 (-2.3) | 0.909 | 0.8 | 0.213 |
|  | Medical doctor | 14 | 7.57±1.01 | 6.23±2.95 | 1.34±2.09 (21.5) | 0.085 | 0.4 | 0.132 |
|  | Physician/Medical Assistant | 19 | 6.58±1.47 | 5.29±2.76 | 1.29±2.43 (24.4) | 0.032* | 0.5 | 0.037* |
|  | CHN/Enrolled nurse | 48 | 5.56±1.38 | 5.02±2.52 | 0.54±2.69 (10.8) | 0.169 | 0.2 | 0.317 |
|  | Pharmacy Technician | 4 | 5.75±0.96 | 7.39±0.68 | -1.64±1.27 (-22.2) | 0.082 | -0.2 | 0.826 |
|  | Midwife | 14 | 4.86±1.99 | 5.58±3.12 | -0.73±2.45 (-12.9) | 0.296 | 0.6 | 0.018* |
| Professional group | Nursing/Midwifery | 140 | 6.00±1.79 | 4.63±2.39 | 1.37±2.59 (29.6) | <0.001* | 0.3 | 0.002* |
|  | Pharmacy staff | 8 | 5.88±2.03 | 6.76±2.44 | -0.88±1.84 (-13.0) | 0.215 | 0.7 | 0.065 |
|  | Prescribers | 33 | 7.00±1.37 | 5.69±2.84 | 1.31±2.50 (23.0) | 0.005* | 0.5 | 0.005* |
| Ever managed snakebite? | No | 29 | 3.97±1.57 | 3.34±2.42 | 0.63±2.19 (18.9) | 0.134 | 0.5 | 0.011* |
|  | Yes | 152 | 6.61±1.47 | 5.23±2.44 | 1.38±2.63 (26.4) | <0.001* | 0.2 | 0.042* |
| Level of confidence of managing snakebite | Not confident | 11 | 2.91±1.76 | 2.73±2.36 | 0.18±2.14 (6.6) | 0.784 | 0.5 | 0.124 |
|  | Fairly confident | 79 | 5.21±1.08 | 4.34±2.35 | 0.87±2.41 (20.0) | 0.002* | 0.2 | 0.126 |
|  | Confident | 78 | 7.17±1.00 | 5.66±2.58 | 1.51±2.78 (26.7) | <0.001* | -0.01 | 0.907 |
|  | Very confident | 13 | 8.81±0.69 | 5.77±1.39 | 3.04±1.65 (52.7) | <0.001* | -0.2 | 0.630 |

NPS = Nominal perceived score.; AKS = Actual Knowledge Score; SD = Standard deviation \* Statistically significant

## DISCUSSION

The outcome of a disease condition depends on several factors including the human beings involved in the process; an assertion supported by the Institute of Medicines’ definition of health care quality as the degree to which health care services for individuals and populations increase the likelihood of desired outcomes and are consistent with current professional knowledge [10]. Snakebites have become an event which claims the lives of between 81,000 and 138,000 persons annually most of whom are poor persons in developing countries involved in agriculture to produce food for their nations and for export to bring foreign exchange to their countries [4]. Although some victims of snake bites seek the services of traditional healers many others seek medical assistance from orthodox health facilities where provision of quality healthcare service can ensure the survival of a snakebite victim or eliminate or reduce any post exposure morbidity [6,11].

Increased productivity had been reported among professionals that are confident about the work they do, which ultimately increases the gratification they derive from the job [12]. Lack of confidence by a healthcare professional can result in feelings of inadequacy, frustration as well as helplessness which can result in increased medical errors which thereby increases the chance of health worker related deformity or death occurring [13, 14]. As much as confidence is needed in the performance of duty, over exaggeration of one’s ability is also detrimental. Since envenomation after a venomous snakebite can quickly affect various body systems and ultimately leading to death, if management of the victims and possible adverse effects of the anti-snake venoms are not executed well by a highly knowledgeable and skills health worker, the prognosis may not be good enough. This study found over exaggeration of knowledge on snakebite management across various sociodemographic classifications when the actual knowledge scores on snakebite management was compared with their presumed level of knowledge before the completion of the knowledge assessment section of the questionnaire. The overall knowledge of health workers on snakebite management in this study was below average [10.60±5.62/22 (48.2%)]. This poor knowledge of health care professionals on snakebite management seem to be same irrespective of the level of development of the health sector of countries. Studies from the United Kingdom and Hong Kong recorded low knowledge on snakebite management which is same in several developing countries such as Loas PDR, Bangladesh, Cameroon, Nigeria [15 – 20]. Males in this study were significantly more knowledgeable about snakebite management than their female counterparts who even significantly exaggerated their knowledge level. The lower knowledge base of females on snakebite management can be attributed to the fear women generally have for snakes [21]. Michael, et al., (2018) did not however find any association between sex of respondents and their knowledge level [20]. It is not clear why the females significantly over exaggerated their knowledge which is in contrast with results of a studies that found men to over exaggerate their capabilities and were also less honest [22, 23]. In this study, health workers in the hospitals were significantly less knowledgeable about snakebite management that colleagues in the lowest level health facility in Ghana, the CHPS zone (p < 0.001) who as well exhibited overestimated confidence of 37.4% compared to the under exaggeration of −25.4% by those working in the CHPS zones. For health workers in the hospitals, there seem to be a moderate but significant correlation between the perceived and actual knowledge on snakebite management (r = 0.5; p < 0.001). The disparity in knowledge levels by the higher and lower level health facility can be due to the more exposed those in CHPS zones are to snakebite issues than those in the hospitals. This results then places snakebite victims that are sent or referred to these higher-level health facilities at higher risk of mismanagement. Among the various health professions, the medical doctors in this study were significantly the most knowledgeable and the nurses least knowledgeable on snakebite management just as reported in some previous similar studies [17, 19]. This is understandable since the medical doctor play the leading role in the management of all cases in the hospitals. The 21.5% over exaggeration of knowledge by the physician and a higher and significant (24.4%; p = 0.037) over estimation of knowledge by the physician/medical assistants can be detrimental to their effective management of cases. The nurses who exhibited the least knowledge level just as in some earlier studies were also the same health professional group that overestimated their knowledge level the most (53.7%, p < 0.001) [17 – 19]. This study found those who had ever managed or ever been trained on snakebite management to be significantly more knowledgeable than those never managed a case or had no previous training (p < 0.001). Effect of training or experience on better management of snakebite had also been observed in some earlier studies in Cameroon, Lao PDR, and Nigeria [17, 19, 20]. Health professionals who had no training but mostly obtained their skills by observing their senior colleagues rather overestimated their knowledge level (38.1% vrs 7.5%). On the other hand, respondents who had ever managed cases although significantly more knowledgeable (p < 0.001), also overestimated their knowledge level (26.4% vrs 18.9%). This over exaggeration of snake management skills for the untrained and even those who had ever managed snakebite cases can adversely affect management of snakebite victims as they will be inappropriately more confident as they even administer or manage such cases wrongly. The effect of high confidence level on the knowledge of respondents was succinctly exhibited when differences between perceived and actual knowledge scores were analyzed. The more confidence a health worker expresses, the higher the over estimation of knowledge; those not confident (6.6%) and very confident (52.7%). Although the overall knowledge on snakebite management may be low, there were some areas where they showed some good knowledge especially those about the 20 minutes whole blood count test, anti-snake venom being the only specific antidote for envenomation and the best route for administering being intravascular. Management of ASV adverse drug reaction (ADR) was rather poorly answered. For more than half (51.6%) of respondents to opt for hydrocortisone rather than adrenaline (22.0%) as their 1^st^ choice in the management of ASV associated adverse drug reaction is a source of worry. This result is even better than a study involving only physicians in a developed country such as Hong Kong, where 57% also opted for hydrocortisone and other antihistamines to manage ASV-induced anaphylactoid reactions [15]. However, up to 90.8% of health workers in the Laos PDR study chose adrenaline as their drug of choice for the management of ASV induced adverse drug reaction [17]. Respondents in this study also exhibited paucity in knowledge on the route of administrating of ASV adverse reaction antidote (19.4%). This poor knowledge on the management of ASV associated ADR seem to be common among health workers across the world as it was reported in India and Hong Kong [15, 24]. ASV associated ADRs are common and known to occurs in between 25% and 62% of victims of snakebite which shows that some morbidity and mortality of snakebites are may not be due to the envenomation only but also mismanagement of the ADR associated with its management [25 – 29].

Results of this study being the first in Ghana we believe should make managers of health systems in Ghana and other developing countries see the need to include snakebite management in the curriculum of their health training institutions. Again, they will formulate policies that will ensure more frequent in-service training on snakebite management for all health workers. Governments should also stock health facilities in rural areas with anti-snake venoms since that was the most stated limitations most health workers indicated as one that affects their facilities ability to manage such cases. For almost half of respondents have managed between 1 and 5 snakebite cases within half a year, shows that snakebite is disease in a rural set up as the Tongu districts. This study however presents some limitations. The study took place in only three out of about two hundred and sixty districts of Ghana so may not represent the situation across the country. Again, generalization of the results may not be appropriate since convenience sampling was used in the selection of the nursing professionals which introduced some biases in the selection of this category of respondents.

## CONCLUSION

There is a deficit in knowledge in the management of snakebite cases among health care professionals in the three Tongu districts of the Volta region with a significant number over estimating their knowledge levels which can lead to mismanagement of victims of snakebite. There is the need for more in-service training on health professionals on snakebite management and should also include issues related to the management of adverse drug reactions associated with anti-snake venoms.

## DATA AVAILABILITY

The data in Microsoft excel and results of analysis in SPSS that were used to support the findings of this study are available from the corresponding author upon request.

## ACKNOWLEDGEMENT

We wish to acknowledge the support of heads of health facilities where the data were collected. We also acknowledge the support given the team of researchers by health workers in the North, Central and South Tongu districts of the Volta region.

## REFERENCES

1. Carter L. Where are there no snakes in the world. 2018. Available from https://www.snakesforpets.com/where-are-there-no-snakes-in-the-world/.

2. Uetz P., Freed, P. & Hošek, J. (eds.) The Reptile Database. 2019. Available from http://www.reptile-database.org.

3. WHO. Global Snakebite burden. 2018. Available from https://apps.who.int/gb/ebwha/pdf_files/WHA71/A71_17-en.pdf.

4. WHO. Venomous snakes distribution. 2010. Available from http://apps.who.int/bloodproducts/snakeantivenoms/database/

5. WHO.. Guidelines for the management of snakebites, 2nd ed. WHO Regional Office for South-East Asia. 2016. Available from https://apps.who.int/iris/handle/10665/249547

6. Majumder D, Sinha A, Bhattacharya SK, Ram R, Dasgupta U, Ram A. Epidemiological profile of snake bite in South 24 Parganas district of West Bengal with focus on underreporting of snake bite deaths. Indian journal of public health. 2014 Jan 1;58(1):17.

7. Sakajiki AM, Ilah GB, Lukman AA, Yakasai AM. Snake bite envenomation seen at a specialist hospital in Zamfara state, North-Western Nigeria. Annals of Tropical Medicine and Public Health. 2017 Mar 1;10(2):391.

8. Punguyire D, Baiden F, Nyuzaghl J, Hultgren A, Berko Y, Brenner S, Soghoian S, Adjei G, Niyogi A, Moresky R. Presentation, management, and outcome of snake-bite in two district hospitals in Ghana. The Pan African medical journal. 2014;19(219).

9. Ghana Statistical Service. Population and Housing Census Summary Results of Final. 2012. Available from http://statsghana.gov.gh/gssmain/storage/img/marqueeupdater/Census2010_Summary_report_of_final_results.pdf.

10. Wolfe A. Institute of Medicine report: crossing the quality chasm: a new health care system for the 21st century. Policy, Politics, & Nursing Practice. 2001 Aug;2(3):233–5.

11. Musah Y, Ameade EP, Attuquayefio DK, Holbech LH. Epidemiology, ecology and human perceptions of snakebites in a savanna community of northern Ghana. PLoS neglected tropical diseases. 2019 Aug 1;13(8):e0007221.

12. Kahr S. Why confidence is important in the healthcare industry. 2019. Available at https://transparency.kununu.com/why-confidence-is-important-in-the-healthcare-industry/

13. Poghosyan L, Clarke SP, Finlayson M, Aiken LH. Nurse burnout and quality of care: Cross-national investigation in six countries. Research in nursing & health. 2010 Aug;33(4):288–98.

14. Cimiotti JP, Aiken LH, Sloane DM, Wu ES. Nurse staffing, burnout, and health care– associated infection. American journal of infection control. 2012 Aug 1;40(6):486–90.

15. Fung HT, Lam SK, Lam KK, Kam CW, Simpson ID. A survey of snakebite management knowledge amongst select physicians in Hong Kong and the implications for snakebite training. Wilderness & environmental medicine. 2009 Dec 1;20(4):364–72.

16. Ahmad Z. A study of the knowledge and attitudes of emergency physicians and plastic surgeons in the management of snakebites. European Journal of Plastic Surgery. 2009 Jun 1;32(3):141–5.

17. Inthanomchanh V, Reyer JA, Blessmen J, Phrasisombath K, Yamamoto E, Hamajima N. Assessment of knowledge about snakebite management amongst healthcare providers in the provincial and two district hospitals in Savannakhet Province, Lao PDR. Nagoya journal of medical science. 2017 Aug;79(3):299.

18. Ahsan HN, Rahman MR, Amin R, Chowdhury EH. Knowledge of Snake bite management among health service providers at a rural Community of Bangladesh. Journal of Current and Advance Medical Research. 2017;4(1):17–22.

19. Taieb F, Dub T, Madec Y, Tondeur L, Chippaux JP, Lebreton M, Medang R, Foute FN, Tchoffo D, Potet J, Alcoba G. Knowledge, attitude and practices of snakebite management amongest health workers in Cameroon: Need for continuous training and capacity building. PLoS neglected tropical diseases. 2018 Oct 25;12(10):e0006716.

20. Michael GC, Grema BA, Aliyu I, Alhaji MA, Lawal TO, Ibrahim H, Fikin AG, Gyaran FS, Kane KN, Thacher TD, Badamasi AK. Knowledge of venomous snakes, snakebite first aid, treatment, and prevention among clinicians in northern Nigeria: a cross-sectional multicentre study. Transactions of The Royal Society of Tropical Medicine and Hygiene. 2018 Feb 1;112(2):47–56.

21. Rakison DH. Does women’s greater fear of snakes and spiders originate in infancy? Evolution and Human Behavior. 2009 Nov 1;30(6):438–44.

22. Byrnes JP, Miller DC, Schafer WD. Gender differences in risk taking: a meta-analysis. Psychological bulletin. 1999 May;125(3):s367.

23. Grosch K, Rau HA. Gender differences in honesty: The role of social value orientation. Journal of Economic Psychology. 2017 Oct 1;62:258–67.

24. Deshpande RP, Motghare VM, Padwal SL, Pore RR, Bhamare CG, Deshmukh VS, Pise HN. Adverse drug reaction profile of anti-snake venom in a rural tertiary care teaching hospital. Journal of Young Pharmacists. 2013 Jun 1;5(2):41–5.

25. Deshmukh VS, Motghare VM, Gajbhiye D, Sv B, Deshpande R, Pise H. Study on acute adverse drug reactions of antisnake venom in a rural tertiary care hospital. Asian journal of pharmaceutical and clinical research. 2014;7(5).

26. Isbister GK, Brown SG, MacDonald E, White J, Currie BJ. Current use of Australian snake antivenoms and frequency of immediate-type hypersensitivity reactions and anaphylaxis. Medical Journal of Australia. 2008 Apr;188(8):473–6.

27. de Silva HA, Ryan NM, de Silva HJ. Adverse reactions to snake antivenom, and their prevention and treatment. British journal of clinical pharmacology. 2016 Mar;81(3):446–52.

28. Shende M, Gawali S, Bhongade K, Bhuskade V, Nandgaonkar A. Studies of adverse drug reaction profile of anti-snake venom at district general hospitaL. Indo American Journal of Pharmaceutical Sciences. 2017 Sep 1;4(9):3033–9.

29. Deva Kumar K. Adverse drug reactions of anti-snake venom among haemotoxic and neurotoxic snake bite: A prospective observational study (Doctoral dissertation, Dhanalakshmi Srinivasan Medical College and Hospital, Perambalur).

